# Discovery and characterization of highly potent and selective covalent inhibitors of SARS-CoV-2 PLpro

**DOI:** 10.1101/2023.05.02.539082

**Authors:** Jian Wu, Tasneem M. Vaid, Hoshin Kim, Jun Lu, Robel Demissie, Hyun Lee, Leslie W.-M. Fung, Jia Chen, Chenghao Xi, Zhezhou Yang, Yu Huang, Zhantao Zhang, Jingqing Zhang, Fengfeng Yan, Michael E. Johnson, Min Li

## Abstract

Coronavirus infections, such as the global COVID-19 pandemic, have had a profound impact on many aspects of our daily life including working style, economy, and the healthcare system. To prevent the rapid viral transmission and speed up recovery from the infection, many academic organizations and industry research labs have conducted extensive research on discovering new therapeutic options for SARS-CoV-2. Among those efforts, RNA-dependent RNA polymerase (RdRp) inhibitors such as Remdesivir, Molnupiravir and 3CLpro inhibitor such as Nirmatrelvir (Paxlovid™) have been widely used as the therapeutic options. Given the recent emergence of several new variants that caused a resurgence of the virus, it would be beneficial to discover more diverse therapeutic options with novel anti-viral mechanisms. In this regard, PLpro has been highlighted since it, along with 3CLpro, is one of the two most important proteases that are required for SARS-CoV-2 viral processing. While 3CLpro inhibitors were extensively investigated in the light of Emergency Use Authorizations of Nirmatrelvir, PLpro inhibitors have not been thoroughly investigated even preclinically. Thus, discovery efforts on antivirals acting against PLpro will be valuable. PLpro inhibitors may exert their activity by inhibiting viral replication and enhancing the host defense system through blocking virus-induced cell signaling events for evading host immune response. In this study, we report the discovery and development of two covalent irreversible PLpro inhibitors, HUP0109 and its deuterated analog DX-027, out of our quest for novel anti-COVID 19 therapeutic agents for the past two and half years. HUP0109 selectively targets the viral catalytic cleft of PLpro and covalently modifies its active site cysteine residue (C111). Promising results from preclinical evaluation suggest that DX-027 can be developed as a potential COVID-19 treatment.

## Introduction

In the past two decades, there have been three major coronavirus outbreaks, which are 2019-nCoV outbreak (SARS-CoV-2, 2019), MERS (2012) and SARS (SARS-CoV, 2002). The most recent coronavirus outbreak, also known as the COVID-19 pandemic, brought huge impacts on our society, especially on medical care, social interactions and organizations, and the global economy. Globally, over 620 million cases of COVID-19 have been reported by February 2023 with 6.8 million deaths.^1^ Although the virus has become more manageable now, through the introduction of vaccines and novel therapies, than the early phase of the outbreak, SARS-CoV-2 is unlikely to be completely eradicated due to several factors including high contagiousness, persistence in animal reservoirs, difficulty in eliminating asymptomatic cases, and constant viral mutations over time. While vaccines have provided significant protection by enhancing immunity against SARS-CoV-2, prophylactic and therapeutic agents targeting SARS-CoV-2 viral replication are needed globally to boost the recovery rate and reduce hospitalizations and deaths. In this regard, Remdesivir (VEKLURY™) has been used and is the only drug that received an NDA approval by FDA. However, delivery of Remdesivir through IV infusion has limited patient compliance and wide application of the therapeutic, as it requires high volume administration and intensive hospital resources. Nirmatrelvir (NTV) and Molnupiravir (MPV) were developed as oral treatments and received Emergency Use Authorizations (EUA) from the FDA (2021). However, MPV showed limited clinical benefits in recovery and prevention of spreading SARS-CoV-2.^2^ Repeated use of the same therapeutic option over time can cause emergence of drug-resistant variants, which in turn requires more diverse therapeutic interventions under different mechanisms of action.

SARS-CoV-2 encodes two distinctive cysteine proteases, namely, the 3-chymotrypsin-like protease (3CLpro, also known as the main protease-Mpro) and the papain-like protease (PLpro). Both 3CLpro and PLpro are essential to the virus proliferation cycle through their involvement in viral protein processing.^3^ Due to the crucial role of 3CLpro in viral life cycles, therapeutic agents targeting 3CLpro have been extensively explored, including Nirmatrelvir,^3c, 4, 12^ while only a few chemical entities targeting PLpro have been reported.^5^ PLpro is responsible for processing three cleavage sites of the viral polyprotein to release mature non-structural proteins 1, 2 and 3. Inhibiting PLpro activity has been shown to be effective in reducing virus replication in cell culture.^5d, 6^ Therefore, PLpro is a promising target of novel antiviral drugs for a range of viral infections, including SARS-CoV-2.

SARS-CoV and SARS-CoV-2 share many common features including pathogenesis, transmission and genetic similarity (∼80%).^7^ Previous studies using SARS-CoV proteases have shown that PLpro is essential for viral protein production and infectivity of coronavirus. Specifically, PLpro is required for viral polyprotein maturation and deconjugates ubiquitin from various substrates involved in maintaining host cell immunity by producing interferon.^8^ Therefore, antiviral drugs targeting PLpro may exert a dual mechanism of action by not only inhibiting viral replication, but also blocking virus-induced cell signaling events that can compromise the host defense.^9^ Since many key features are conserved between SARS-CoV and SARS-CoV-2, the results of previous studies on the structure and function of PLpro in SARS-CoV can inform the development of inhibitors against PLpro of SARS-CoV-2. Based on these similarities, we decided to utilize the published research data on SARS-CoV with naphthalene-based small molecule inhibitors for SARS-CoV-2.^5a, b, d, 9a, 10^

As of Feb 27, 2023, the WHO has identified α-20I, β, γ-20J, δ-21A and ο-21K to 23A as major variants of concern.^1, 11^ In the variants, three PLpro residues are reported as defining mutations: A145D (ORF1a 1708) in α-20I, K232Q (ORF1a 1795) in γ-20J, P77S (ORF1a 1640) in δ-21A and ο-22D. Two additional PLpro mutant residues found in the variants are A246T (ORF1a 1809) in η-21D (count=521; 6.83%) and G256S (ORF1a 1819) found in ο-22F (count=2115; 2.65%) and in ο-23A (count=1086; 2.73%). None of the five residues are conserved in SARS-CoV-2 PLpro. A246 is the only residue close to the channel formed by binding loop 2 (BL2), and is located on a β-strand, oriented away from the binding site, about 8.5 Å distant from the edge of the BL2 channel, but is not found in currently dominant ο-variants. The rest of the mutation sites are more than 10 Å distant from any portion of the binding region, including the catalytic triad.^3b^ Thus, we do not expect PLpro inhibitors to be affected by any of the known mutations found with significant frequency, which makes Plpro as an attractive target for SARS-CoV2 since PLpro inhibitor blocking the binding region near the catalytic triad would still be efficacious.

Previous attempts to design inhibitors against SARS-CoV PLpro yielded some promising results for a family of naphthalene-based inhibitors.^5a, 9a^ GRL0617 was identified as a selective inhibitor of PLpro of SARS-CoV through a HTS screen and optimization by the Ghosh group.^5a-c^ With the SARS-CoV outbreak effectively contained in 2003 with no re-emergence, the developmental efforts on these potential anti-CoV therapeutics were discontinued. Also, several potential issues of these compounds including weak cellular antiviral activity and metabolic liability limited their advancement as an antiviral drug. However, recent preliminary research on small molecule inhibitors of SARS-CoV PLpro have shown their antiviral activities against SARS-CoV-2 PLpro as well (IC_50_ = 2.3 μM),^3b^ which encouraged us to invest our discovery efforts further on these chemical scaffolds for novel antivirals against SARS-CoV-2.

## Results and Discussion

### Identification of HUP0109 as the new lead candidate

Since the catalytic sites of PLpro between SARS-CoV and SARS-CoV-2 share significant similarity and GRL0617 has shown moderate potency in vitro against PLpro of SARS-CoV-2, we designed compounds based on GRL0617 and its analogs. Through systematic analysis of available medicinal chemistry results on SARS-CoV and diverse structure classifications of GRL0617 analogs, we chose two main chemical scaffolds from the naphthalene series that include aniline or piperidine as the pharmacophore. To improve efficiency and accuracy in the design of analogs, we employed virtual screening using the ICM Molsoft docking method based on a 4-dimensional docking approach utilizing a crystal structure with non-covalent inhibitor GRL0617 (PDB ID: 7JRN) and two crystal structures with peptidomimetic covalent inhibitors, VIR251 and VIR250 (6WX4 and 6WUU). The docking calculations were followed by molecular dynamics simulations to calculate binding free energies and gauge the conformational stability of the docked compounds. Several hundred analogs were designed to target the binding site of the catalytic cleft of PLpro. To improve the binding affinity mechanistically based on the preliminary results of peptidomimetic analogs^10e^ utilizing binding mode of LRGG of AMC PLpro substrate, we designed covalent compounds targeting the active cysteine (C111) in the catalytic site by introducing various combinations of diverse linkers and covalent warheads to extend the compound. Then we performed a comprehensive medicinal chemistry campaign to yield potent compounds possessing potentially better drug-like properties. The enzymatic activity was evaluated using the Z-RLRGG-AMC assay.^13^ Through systematic SAR studies using the PLpro enzymatic assay, we produced a series of covalent inhibitor analogs that were biochemically more potent than GRL0617 (> 50 analogs with IC_50_ > 10-fold improvement). Encouraged by the significant improvement of potency at the enzymatic level, a handful of compounds were further tested in cellular antiviral activity assay using A549:hACE2 cells infected by several strains including the original Washington strain, delta and omicron variants. Using these biochemical and cellular evaluations, HUP0109 was identified as our first lead PLpro inhibitor with excellent potency and selectivity (Figure 1). HUP0109 inhibited PLpro with an enzymatic IC_50_ of 0.02 μM, indicating a >100-fold increase in potency compared to GRL0617. Potency improvement at the cellular level was further confirmed by several compounds (>10 compounds, >10-fold). Of note, the improvement of biochemical potency of HUP0109 translated very well into the cellular antiviral activity (EC_50_ = 0.05 µM, >100-fold compared to GRL0617), as shown in Figure 1C. In addition, the exquisite selectivity of HUP0109 for PLpro of SARS-CoV-2 was demonstrated by little or no inhibitory effect of HUP0109 on other deubiquitinases or cysteine proteases, including USP7, USP2, USP18, UCHL1, UCHL3, Caspase3, Cathepsin B, and Cathepsin L. No cytotoxicity was observed in the tested concentrations (up to 4 µM) in the A549:hACE2 cell line.

**Figure 1.**
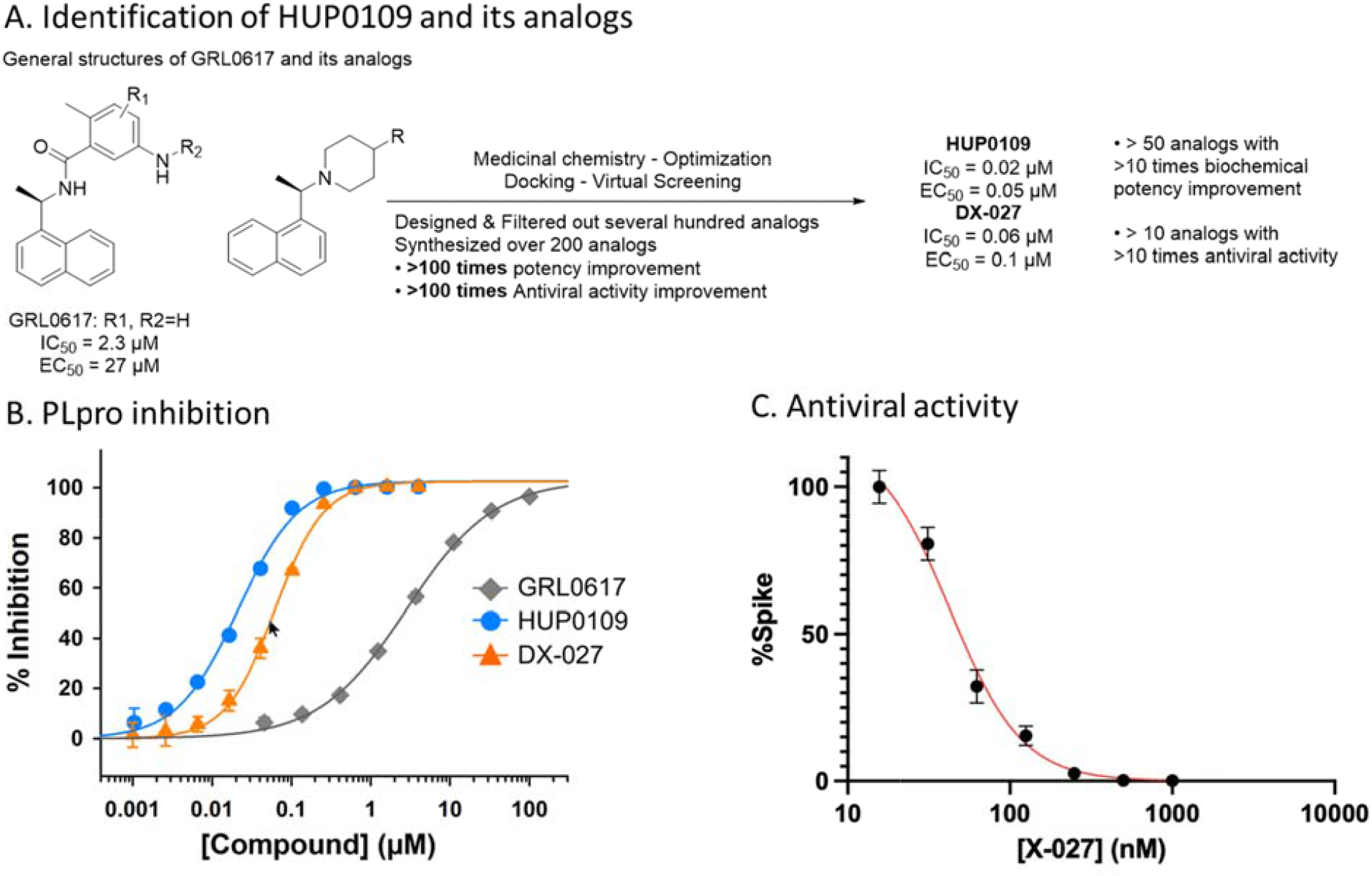
Summary of identification of HUP0109 and its analogs. A. Identification of HUP0109 and its analogs through medicinal chemistry campaign starting from GRL0617 and its analogs. B. PLpro enzymatic inhibition plot of GRL0617, HUP0109 and DX-027. C. Antiviral activity plot of X-027 in A549:hACE2 cells infected by the Omicron B.1.1.529 strain. The EC_50_ was determined by measuring the % Spike protein as a function of inhibitor concentration. X-027 is the racemic form of HUP0109, while DX-027 is the deuterated analog of the racemic form of HUP0109.

### Covalent mode of inhibition of PLpro by HUP0109

To identify the precise nature of inhibition, WT-PLpro was incubated with HUP0109 and analyzed by MALDI-TOF MS. Interestingly, an increase in mass of ∼528 Daltons (corresponding to molecular weight of HUP0109) was detected when the WT-PLpro had been incubated with HUP0109, indicating the formation of a covalent complex between WT-PLpro and HUP0109 (data not shown). PLpro contains 11 cysteine residues of which one cysteine (C111) is a part of the catalytic triad and four cysteines (C189, C192, C224, C226) are present in the finger region, coordinating to a tightly bound Zn^2+^ ion.^13^ Among the remaining six cysteines (C148, C155, C181, C260, C270, C284), only C270 is surface exposed and could potentially be modified by covalent small molecules. To confirm whether the active site catalytic cysteine (C111) is the only residue targeted by HUP0109, we utilized an active site cysteine to alanine mutant form of PLpro (PLpro C111A). Mutant PLpro protein was incubated with HUP0109 and analyzed by MALDI-TOF MS. Remarkably, replacement of the active site cysteine with alanine completely blocked the formation of compound adduct on the PLpro C111A mutant, confirming that the active site cysteine is indeed the only conjugation site for HUP0109 under these conditions. The closely related analog of HUP0109, in which the covalent warhead was replaced by carboxylic ester, was devoid of activity, suggesting the importance of covalency in this chemical series. In addition, X-027 (racemic HUP0109) inhibits PLpro with an enzyme inactivation rate (k_inact_) to inhibition constant (K_i_) ratio (k_inact_/K_i_) of 2664 ± 698 M^−1^s^−1^, where k_inact_/K_i_ is a second-order rate constant describing the efficiency of the overall conversion of free enzyme to the covalent enzyme-inhibitor complex.

### Synergetic potential using HUP0109

There are potential benefits of combining PLpro inhibitor with other mechanism-based antivirals since 3CLpro and PLpro are the two main enzymes responsible for the replication of the SARS-CoV-2. The pilot combination study of HUP0109 analogs and Nirmatrelvir suggested promising synergistic activity especially at the concentrations below 30 nM (data not shown), which warrants further investigations to find out the best combination of PLpro and other mechanistic antivirals. It is important to characterize the risk of drug-drug interactions (DDIs) early in the drug discovery process, especially when the potential for synergy is a critical consideration. Of note, X-027 did not inhibit transporters (BCRP and P-gp) indicating a low risk of transporter-mediated DDIs.^14^

### Favorable preclinical ADME and pharmacokinetic profile of X-027 (racemic HUP0109)

The pharmacokinetics of X-027 was characterized both in vitro and in vivo. The hepatocyte metabolic stability study suggested that X-027 has moderate CL_int_ in mouse (167 ml/min/kg), rat (72.5 ml/min/kg), dog (65 ml/min/kg) and human (18 ml/min/kg) (Table 1). X-027 metabolism was significantly inhibited (≥70%) by a potent CYP3A4/5 inhibitor ketoconazole, which suggests that X-027 is mainly metabolized by CYP3A4/5. The predicted major metabolism was the hydroxylation nearby the covalent warhead area of X-027. In light of the recent success of Nirmatrelvir combination with ritonavir which improved the PK profile of Nirmatrelvir in the clinic, the in vivo PK of X-027 was profiled in mice with ritonavir. As expected, there was a significant increase in X-027’s plasma exposure (>10-fold) by blocking the CYP3A4 metabolism via ritonavir (Table 2).

**Table 1.** In vitro metabolic stability of X-027 in hepatocytes from preclinical species and human

| Compound | Hepatocytes | $t_{1/2}$ (min) | In vitro $CL_{int}$ (mL/min/kg) |
| --- | --- | --- | --- |
| X-027 | Mouse | 98 | 167 |
|  | Rat | 89.6 | 72.5 |
|  | Dog | 147 | 65 |
|  | Human | 216 | 18 |

**Table 2.** Single dose pharmacokinetics of X-027

| Compound | Species | Dose (mg/kg) | $T_{1/2}$ (h) | $C_{max}$ (ng/mL) | $AUC_{last}$ (ng·h/mL) | MRT (h) |
| --- | --- | --- | --- | --- | --- | --- |
| X-027 | mouse | 25 | 1.11 | 278 | 426 | 1.44 |
|  |  | 25+ritonavir | 3.39 | 1810 | 4661 | 2.61 |
|  | rat | 50 | 3.80 | 320 | 1278 | 4.65 |
|  | dog | 50 | 2.63 | 2130 | 8211 | 3.94 |

Since the CL_int_ in rat hepatocytes of X-027 was moderate, we further characterized its oral PK in SD rats. The oral administration of X-027 at 50 mg/kg showed moderate half-life (T_1/2_ = 3.8 h) and plasma exposure (C_max_ = 320 ng/mL (660 nM), AUC_last_ = 1278 ng·h/mL), which can cover the plasma exposure above EC_90_ (83 nM, 95% CI 75 to 92 nM) over 6 hours (Table 2 and Figure 2). Overall, the compound exhibited decent plasma exposure in oral administration in rats indicating its development potential as an orally available PLpro inhibitor targeting SARS-CoV-2.

**Figure 2.**
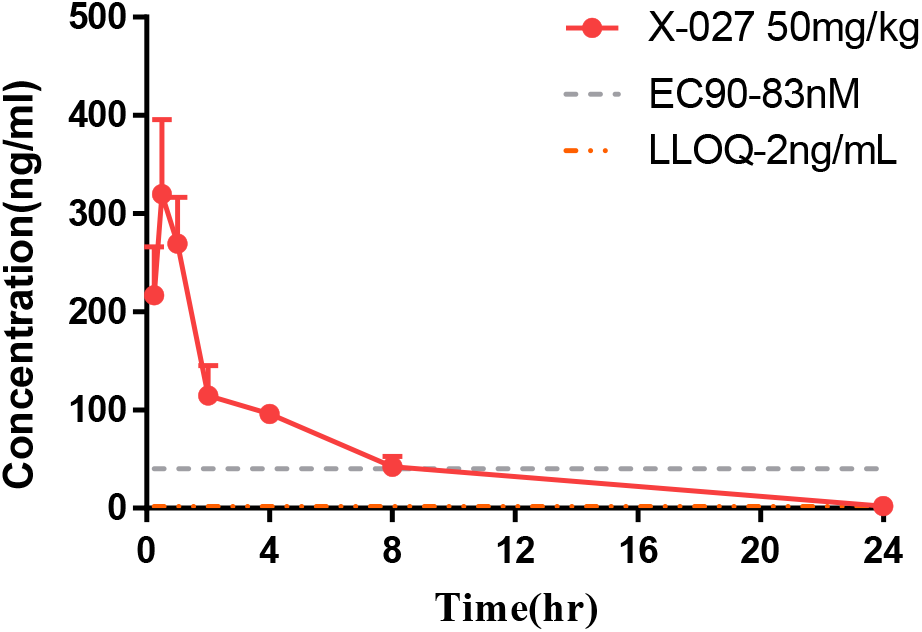
Rat oral exposures of once-daily administered X-027 compared with EC90 value

### Preclinical safety profile supports further development potential of DX-027

Since sufficient exposures were achieved at the low dose in rats (50 mg/kg, PO), we further characterized the in vivo PK of X-027 at higher doses in rats and dogs. In single dose oral PK studies (300 to 2000 mg/kg in rats and 50 to 1000 mg/kg in dogs, Table 3A), X-027 showed no significant adverse effects other than minor GI-related events such as loose stools, and soft stool. In rats, oral administration of X-027 from 100 to 600 mg/kg exhibited more than dose proportional increase in exposure, where AUC increased from 4433 ng·h/mL to 59103 ng·h/mL. Meanwhile, the exposure from 600 to 1000 mg/kg appears to be linear (59103 to 125474 ng·h/mL). No accumulation was noted after the 7-day repeated dosing (Table 3B). The no observed adverse effect level (NOAEL) was 600 mg/kg. In 7-day repeated 1000mg/kg dosing, no significant adverse effects were observed, except minor weight loss (6-10 %) and laboratory abnormalities such as high cholesterol, and low triglycerides.

**Table 3A.** Single PO toxicokinetic studies of X-027

| Compound | Species | Dose<br>(mg/kg) | T<sub>1/2</sub><br>(h) | C<sub>max</sub><br>(ng/mL) | AUC<sub>last</sub><br>(ng·h/mL) | MRT<br>(h) |
| --- | --- | --- | --- | --- | --- | --- |
| X-027 | rat | 300 | 6.3 | 2501 | 34935 | 11.7 |
|  |  | 600 | 177 | 3150 | 56970 | 257 |
|  |  | 1200 | 15.6 | 11557 | 177372 | 24.3 |
|  |  | 2000 | 26.3 | 11848 | 212193 | 39.7 |
|  | dog | 50 | 2.6 | 2130 | 8211 | 3.9 |
|  |  | 200 | 6.2 | 3270 | 19579 | 6.7 |
|  |  | 400 | 22.3 | 7465 | 54478 | 7.3 |
|  |  | 1000 | 12.7 | 9355 | 74813 | 8.3 |

**Table 3B.** 7-day oral PK of X-027 in rats

| Compound | Day | Dose<br>(mg/kg) | C<sub>max</sub><br>(ng/mL) | AUC<sub>last</sub><br>(ng·h/mL) |
| --- | --- | --- | --- | --- |
| X-027 | 1 | 100 | 616 | 4433 |
|  |  | 600 | 3393 | 59103 |
|  |  | 1000 | 9172 | 125474 |
|  | 7 | 100 | 616 | 3695 |
|  |  | 600 | 3332 | 30442 |
|  |  | 1000 | 10303 | 103733 |

To further improve the PK of X-027, we embarked on another medicinal chemistry campaign to block major metabolism by deuterating the potential site of hydroxylation. After a series of chemical modifications, the deuterated analog DX-027 was successfully produced that exhibited better PK profile in vivo without compromising activity (IC_50_ = 0.06 µM, EC_50_ = 0.1 µM). Oral administration of DX-027 from 20 to 40 mg/kg in beagle dogs (Table 4) exhibited more than dose proportional increase in exposure where AUC increased from 5229 to 31561 ng·h/mL. Meanwhile, the exposure from 40 to 80 mg/kg dose studies appeared to be linear (31561 to 60055 ng·h/mL). Altogether, DX-027 achieved approximately 4-fold higher exposure in C_max_ and AUC at lower dose (40 mg/kg) than that of X-027 at 50 mg/kg. In addition, the concentration of X-027 or DX-027 in the lung was approximately five times higher than that of plasma, suggesting that the compound is being distributed to and preferentially accumulated in the lungs, which could lead to better efficacy in the target tissue of interest (Table 5). In vivo PK profiling of X-027 and DX-027 clearly shows the developmental potential of PLpro inhibitors.

**Table 4.** 14-day oral PK of DX-027 in dogs

| Compound | Day | Dose<br>(mg/kg) | C<sub>max</sub><br>(ng/mL) | AUC<sub>last</sub><br>(ng·h/mL) |
| --- | --- | --- | --- | --- |
| DX-027 | 1 | 20 | 2390 | 5229 |
|  |  | 40 | 9040 | 31561 |
|  |  | 80 | 17800 | 60055 |
|  | 14 | 20 | 2505 | 5653 |
|  |  | 40 | 12850 | 49488 |
|  |  | 80 | 23000 | 101586 |

**Table 5.** Target tissue distribution in rats and dogs

| Compound | X-027 | DX-027 |
| --- | --- | --- |
| Species | Rats | Dogs |
| Dose (mg/kg) | 50 | 50 |
| Lung/Plasma* | 5.9 | 5.5 |

**\*Note:** Lung/plasma concentration ratio at T<sub>max</sub> after single oral administration in rats and dogs.

Compelling data accumulated over COVID-19 pandemic have established PLpro as an antiviral target. In moving from the stage of developing excellent tool compounds such as GRL0617 to that of developing antiviral drugs, however, one of the major challenges lies in developing potent and selective PLpro inhibitors with drug-like properties. Structure guided design and development will be crucial to generate selective PLpro inhibitors. Although PLpro inhibitors have shown antiviral cellular efficacy in a variety of cell lines, very limited in vitro ADME and no in vivo pharmacokinetics data have been presented in the literature for these inhibitors. The availability of PLpro inhibitors with drug-like and well-defined pharmacokinetics properties will be essential for making progress in the therapeutic arena. These properties, however, should be considered in the context of the mechanism of inhibition. In comparison with reversible inhibitors, selective covalent irreversible PLpro inhibitors offer potential advantages. For example, a relatively short duration of treatment could result in sustained PLpro inhibition and prolonged biological effects, leading to improved therapeutic efficacy. Moreover, development of covalent PLpro inhibitors may be governed by pharmacokinetics parameters that differ from those for reversible inhibitors, particularly with respect to half-life and clearance.

Through the medicinal chemistry campaign, we have successfully identified a lead candidate HUP0109 that selectively and potently inhibits PLpro (IC_50_ = 0.02 μM, EC_50_ = 0.05 μM). The covalency of the compound with catalytic Cys in PLpro of SARS-CoV-2 was confirmed. We demonstrated its strong antiviral activity against SARS-CoV-2 in A549:hACE2 cells, and favorable ADME/PK properties of HUP0109 in vivo. We also suggested the potential combination therapeutic approaches with current medications. In addition, we showed its further optimization potential in PK properties with deuterated analog DX-027. Based on the encouraging preliminary in vivo observations at low dose via oral administration, HUP0109 and its analogs are currently under investigation in transgenic mice model of SARS-CoV-2 (K18-hACE2) using delta/omicron variants and we plan to report in vivo efficacy data including virus titer in the lung and survival score. Altogether, the in vitro and in vivo data of HUP0109 and its deuterated analog DX-027 suggest the strong development potential of the novel orally available PLpro inhibitors.

## Methods

### Computational docking and simulation

The docking was performed using ICM Molsoft v3.9-2c GUI.^15^ The 4-dimensional docking approach was utilized using three crystal structures: 7JRN (Crystal structure of the wt SARS-CoV-2 PLpro with inhibitor GRL0617), 6WX4 (Crystal structure of the SARS-CoV-2 PLpro in complex with covalent peptide inhibitor VIR251) and 6WUU (Crystal structure of the SARS-CoV-2 PLpro in complex with covalent peptide inhibitor VIR250). Each ligand was assigned the Merck molecular force field (MMFF)^16^ force field atom types and charges and was then subjected to Cartesian minimization. Two types of grid maps were generated including the residues within 5 Å of the GRL0617 binding pocket (i) Non-covalent maps for the docking of GRL0617 (as a control) and (ii) Covalent maps by selecting C111 as a targeted nucleophile. The Monte Carlo (MC) docking runs were performed three times using an MC thoroughness setting of 10, which controls the length of the run, and the top (i.e. lowest) scoring poses were analyzed. Two types of scores (ICM dock score and RTCNN score) were analyzed to rank the compounds.

The Molecular Dynamics (MD) simulation study was conducted on GPU (NVIDIA RTX 2070) accelerated Gromacs 2019.6 software.^17^ The CGenFF tool was used to generate ligand topology and parameter.^18^ For the protein molecule, the CHARMM36^19^ all-atom force field was used. The protein-inhibitor complex was embedded in a dodecahedron simulation box set to the periodic boundary conditions in three spatial dimensions, with minimum distance being 1 nm. The SPC water model was used and the system was neutralized by adding Na+/Cl–ions. Eventually, the ionic concentration of the solution was set to reach 0.15 M. The production MD was conducted with a time step of 2 fs under constant pressure of 1 atm and constant temperature of 300 K. The steepest descent algorithm of 5000 steps was used to equilibrate and minimize the system. The cut off was set to 1.2 nm for both long-range van der Waals and electrostatic interactions. The long-range electrostatic interactions were modelled using PME.^20^ For binding energy calculations, 2 ns MD simulations were performed, while for the binding conformational stability analysis of the lead compound, long-scale 1 μs simulation was performed. The resulting trajectory was analyzed using VMD.^21^

### PLpro preparation and purification

The PLpro (pp1ab 1564-1878) was codon-optimized, synthesized, and cloned into pET11a vector with a TEV cleavable his6-tag at the N-terminus. The recombinant plasmid was transformed into BL21(DE3) expression cells and grown in Luria– Bertani (LB) media with carbenicillin (100 μg/mL) at 37°C while shaking at 220 rpm until the OD_600_ reached 0.6, when it was induced with 0.5 mM IPTG and incubated for an additional 16 h at 18 °C before harvesting. The cell pellet was resuspended and lysed by sonication in lysis buffer (50 mM Tris, pH 8.0, 500 mM NaCl, 20 mM imidazole, 5 mM β-MCE, 1 mg/mL lysozyme, 1% Triton X-100 and 0.025 mg/mL DNase I). A HisTrap HP column was used to purify the histidine-tagged protein using a stepwise gradient of elution buffer (50 mM Tris, pH 8.0, 500 mM NaCl, 500 mM imidazole and 5 mM β-MCE) with an AKTA Pure FPLC system. The histidine-TEV tag was removed by incubating the eluted protein with 1 unit/100 μg protein of TEV protease at 4 °C for 16 h. The digested protein was reloaded onto a HisTrap HP column equilibrated with 50 mM Tris, pH 8.0, 500 mM NaCl and 5mM β-MCE, and the histidine-TEV tag cleaved PLpro was collected in the flowthrough and loaded onto a HiLoad 16/60 Superdex 75 PG gel filtration column that was equilibrated with 50 mM Tris, pH 8.0, 200 mM NaCl and 1 mM TCEP. Protein samples were analyzed by SDS-PAGE, and the final purity was above 95%.

### PLpro inhibition *In vitro*

The PLpro enzyme was purified as described above and prepared in assay buffer (50 mM HEPES, pH 7.5, 0.01% Triton X-100 (v/v), 0.1 mg/mL BSA, and 2 mM DTT (or 1 mM GSH for covalent inhibitors). IC_50_ values were measured in triplicate. A series of increasing concentrations (0–100 µM final concentration at 2.5-fold serial dilution) in 100% DMSO were prepared in a 384-well plate. 7 µL of 225 nM (3X) enzyme solution was distributed into wells, and 7 µL of varying concentration of 3X compounds were added and incubated for 10 min and 60 min for non-covalent inhibitors and covalent inhibitors, respectively. The enzyme reaction was initiated by adding 7 µL of the 75 µM (3X) substrate, Z-Arg-Leu-Arg-Gly-Gly-AMC (Bachem Bioscience), and its activity was continuously monitored at 360 nm (excitation) and 450 nm (emission) for at least 10 min with BMG Biotech Optima microplate reader. The IC_50_ values were calculated by fitting with the Hill equation (1), with Sigmaplot v14.5, where *y* is percent inhibition, *x* is inhibitor concentration, *n* is the slope of the concentration–response curve (Hill slope), and *V*_max_ is maximal inhibition from three independent assays.

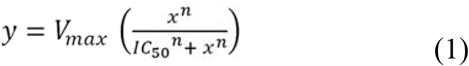

### SARS-CoV-2 antiviral activity Assay

PLpro inhibitor antiviral activities were tested in a cell culture assay with A549:hACE2 cells (Invivogen) by the following protocol: seeded 10K cells (A549:hACE2) per well the day before testing. A serial dilution of the drug in 2% DMEM media was made to add to the cells. Cells were left to incubate with drugs for 2 h prior to infection. Then cells were infected with 0.5 MOI (multiplicity of infection) and 48 hours later they were fixed with 10% Formalin for 15-30 min. The cells then underwent immunohistochemistry antibody staining. Cells were blocked with 1% BSA+.025% Saponin for 1h, washed with 3% hydrogen peroxide for 5 min, washed with PBS and PBST, then primary antibody [mouse anti-Spike antibody (GTX632604, GeneTex)] in blocking buffer overnight. This was followed the next day by additional PBS/PBST washing, secondary HRP antibody for 1 hour then DAB stain for 15 minutes. The percentage of cells positive for spike protein was then assessed under the microscope. The EC_50_ was determined from the following equation (2), where [I] is the inhibitor concentration and n is the Hill coefficient, using Prism (Graphpad Software, San Diego). Compounds were tested against the Washington and Omicron B.1.1.529 and B.5 variants. Cytotoxicity was tested in the same cell line at concentrations up to 4 µM. Synergy was calculated using the BLISS algorithm in the SynergyFinder R web server.^22^

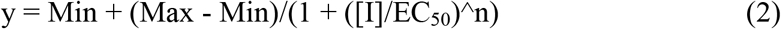

### Compound binding assay and inactivation rate assay

The purified SARS-CoV-2 PLpro enzyme was immobilized on flow channels 2 and 4 of a CM5 sensor chip using standard amine-coupling with running buffer HBS-P (10 mM HEPES, 150 mM NaCl, 0.05% surfactant P-20, pH 7.4) using a Biacore T200 instrument. Flow channels 1 and 3 were used as control surfaces. The PLpro enzyme was diluted in 10 mM sodium acetate (pH 5.0), and immobilized after sensor surface activation with 1-ethyl-3-(3-dimethylaminopropyl) carbodiimide hydrochloride (EDC)/N-hydroxy succinimide (NHS) mixture followed by ethanolamine (pH 8.5) blocking on unoccupied surface area. Selected compounds were initially prepared as 10 mM DMSO stock solutions, and compound solutions with a series of increasing concentrations (0 – 10 µM at 4-fold dilution) were applied to all four channels at a 30 µL/min flow rate at 25 °C. The single-cycle kinetic method was run, and real-time response units were monitored. Sensorgrams were double referenced with the blank channel and zero concentration responses and fitted with 1 to 1 Langmuir kinetic equation embedded in the Biacore Insight software. Surface Plasmon Resonance (SPR) binding analysis showed the direct binding response of selected PLpro inhibitors to the immobilized SARS-CoV-2 PLpro protein at a series of increasing concentration using single-cycle kinetics. K_d_ values were determined for representative compounds.

The purified SARS-CoV-2-PLpro protein was prepared in the assay buffer (50 mM HEPES, pH 7.5, 0.01% Triton X-100, 0.1 mg/mL BSA). The PLpro substrate, Z-RLRGG-AMC (Bachem Bioscience), is a small fluorogenic peptide. The cleavage of the substrate by PLpro releases the AMC (7-amido-4-methylcoumarin) which generates fluorescence signal. Assays were done using black low-volume 384-well plates (Greiner). All compounds were initially prepared as 1 mM stocks in 100% DMSO. A series of increasing concentrations were prepared in 100% DMSO through 2-fold serial dilutions. Final compound concentrations (1x) tested ranged from 1.5-1500 nM. 7 µL of the compound (3x) solutions and 7 µL of the substrate (3x) solution were added to the wells. The reaction was initiated by adding and mixing 7 µL of the SARS-CoV-2-PLpro (3x) solution. The final concentration (1x) of the SARS-CoV-2-PLpro was 20 nM and the final concentration (1x) of the substrate was 10 µM. The fluorescence intensity at 450 nm (excitation at 360 nm) was continuously monitored for 1.5 hr at 30 °C using the BMG LabTech PolarStar Optima Plate Reader. Using DynaFit, the kinetic curves were fit to the equation: F = F_0_ + r_P_[S]_0_[1-exp[-β(1-exp(-αt))]] to derive the values for α parameters. K_inact_/K_i_ was derived by fitting a hyperbolic equation, α=K_inact_([I]_0_/([I]_0_ + K_i_)), to the plot of α versus compound concentration.^23^

### Covalent mode of inhibition determined by MALDI-TOF MS

Covalent binding and selectivity for the catalytic cysteine was determined by comparing MALDI mass spectrometric analyses of wild-type SARS-CoV-2 PLpro incubated with inhibitor with that of C111S-CoV-2 PLpro, in which the catalytic cysteine was mutated to serine. Both WT- and C11S-PLpro proteins were prepared at 10 µM in 50 mM HEPES buffer at pH 7.5 and incubated with either DMSO or 5-fold higher concentration (50 µM) of the tested compounds for 2 hours at room temperature. Zip-Tip(C4) was performed on the samples prior to the analysis to remove the salt. The samples were aspirated and dispensed through Zip-Tip to bind, wash, and elute. Recovered samples were salt-free and eluted in 4 μL of 50% Acetonitrile with 0.1% trifluoroacetic acid. The matrix-assisted laser desorption/ionization time-of-flight mass spectrometry (MALDI-TOF MS) measurements were performed with a Bruker MALDI TOF Microflex (Bruker, Bremen, Germany) instrument equipped with 60 Hz N2-Cartridge-Laser with maximum repetition rate of 20 Hz, capable of executing both linear and reflector modes. Mass spectra were acquired in the linear positive ion mode by summing spectra from 500 random laser shots at an acquisition rate of 60 Hz. Treatment of PLpro with representative compounds resulted in a covalent irreversible adduct formation.

### In vitro metabolic stability in hepatocytes from preclinical species and human

Metabolic stability was examined using 1 μM of the tested compounds and hepatocytes 1.0 × 10^6^ cells/mL. Incubations were conducted in duplicate. At 0, 15, 30, 60, 90 and 120 min, aliquots were removed and added to acetonitrile containing internal standard to terminate the reaction. Protein was removed by centrifugation and the supernatant was analyzed by ultra-performance liquid chromatography-tandem mass spectrometry (UPLC-MS). Mean stability data (half-life (t_1/2_) and intrinsic clearance (CL_int_)) is depicted.

### Reaction phenotyping of X-027 metabolism

Metabolic stability of X-027 (10 μM) was examined in HLM (human liver microsomes, 1 mg/mL protein concentration) with potassium phosphate (100 mM, pH 7.5) containing MgCl_2_ (3.3 mM) and NADPH (10 mM). To probe the reaction phenotyping of X-027 metabolism, selective CYP1A2 inhibitor naphthoflavone, CYP2C9 inhibitor sulfaphenazole, CYP2C19 inhibitor benzylnirvanol, CYP2D6 inhibitor quinidine, CYP3A4/5 inhibitor ketoconazole were also included to the incubations. At 5, 10, 15, 30, 60 min, aliquots were removed and added to acetonitrile containing internal standard to terminate the reaction. Protein was removed by centrifugation and the supernatant was analyzed by ultra-performance liquid chromatography-tandem mass spectrometry (UPLC-MS).

### Plasma sample analysis

Concentrations of tested compounds in the plasma samples were analyzed using LC-MS/MS method. Shimadzu HPLC LC-30AD and Triple Quad 5500 with Analyst software 1.6.3. were used. Samples were injected onto a column (Waters ACQUITY UPLC® BEH C18 1.7μm (2.1×50mm)) maintained at 40 °C, at a flow rate of 0.6 mL/min. The mobile phase components were A: 0.1% formic acid in water and B: 0.1% formic acid in acetonitrile. The initial mobile phase composition of 5% B was held for 0.5 min followed by sequential linear gradients to 95% B at 1.3 min. This composition was held for 0.5 min followed by re-equilibration at initial conditions for 0.5 min.

### In vivo efficacy studies

Appropriate numbers of B6.Cg-Tg (K18-ACE2)2 mice (Jackson Lab) were ordered, acclimated and divided into several groups (cohorts) with equal numbers in each group. On Day 0, each mouse was weighed, anesthetized by IP injection of ketamine (50-60 mg/kg) and xylazine (3-6 mg/kg) and challenged intranasally with SARS-CoV-2 (Delta) (20,000 PFU). Treatment with test compounds was followed after 6 hours. Mice in Cohort 1 were used as negative control, with p.o. vehicle only, by oral gavage for each mouse per dosing. Mice in Cohort 2 were used as positive control, with Nirmatrelvir (300 mg/kg), oral gavage for each mouse per dosing. Mice in other Cohorts were treated with the compounds, by oral gavage. Days 1-4, the same treatments, twice daily (6 am and 6 pm), were applied to the mice in each Cohort. On Day 5, only one treatment was applied to mice in each Cohort at 6 am. All mice were then euthanized, and lungs were removed, with one lung of each mouse fixed in 10% formalin and the remaining lung divided into 2 sections. One section was placed into 500 µL DMEM, and the other section into 500 µL RNA buffer. The sections were flash frozen on dry ice and stored at - -80 °C. The fixed lung samples were removed from tubes and placed in fresh 10% formalin, validated for viral inactivation, placed into cassettes and sent to histology lab for sectioning, slide preparation and subsequent digitization for pathology review. Lung samples in DMEN were thawed, homogenized, and tested for virus quantification, using TCD150 assay. Lung samples in RNA buffer were thawed, homogenized, and tested for virus quantification, using RT-qPCR assay.

## Acknowledgment

We thank the staff of the Howard Taylor Ricketts Laboratory, University of Chicago, Lemont, IL for performance of the antiviral EC50 measurements. The technical assistance of Mr. Hanbin Shan during the scale up synthesis of several key compounds is highly appreciated.

